# Decomposing virulence to understand bacterial clearance in persistent infections

**DOI:** 10.1101/2021.03.29.437521

**Authors:** Beatriz Acuña Hidalgo, Luís M. Silva, Mathias Franz, Roland R. Regoes, Sophie A.O. Armitage

## Abstract

Hosts are not always successful at controlling and eliminating a pathogen and the factors causing variation in pathogen clearance are not well understood. To address this problem, we used *Drosophila melanogaster* to investigate how infections with different bacterial pathogens affects virulence, clearance and persistence. In this context we developed novel hypotheses that focus on how variation in clearance should be related to variation in different components of virulence, where virulence is the infection-related reduction in host fitness. To achieve this, virulence was decomposed into exploitation, i.e., how well bacteria can replicate inside the host, and per parasite pathogenicity (PPP), i.e., the amount of damage per parasite inflicted on the host. We used four bacterial species: *Enterobacter cloacae*, *Providencia burhodogranariea, Lactococcus lactis* and *Pseudomonas entomophila*. The injection doses spanned four orders of magnitude, and survival was followed to estimate virulence. Bacterial load was quantified in live flies during the acute (1-4 days) and chronic (7-35 days) phases of infection, and we tested infection status of flies that had died up to ten weeks post infection. We show that sustained persistent infection and clearance are both possible outcomes for bacterial species across a range of virulence. Bacteria of all species could persist inside the host for at least 75 days, and injection dose partly predicted within-species variation in clearance. Our decomposition of virulence showed that species differences in bacterial virulence could be explained by a combination of variation in both exploitation and PPP. In addition, we found that that higher exploitation leads to lower bacterial clearance, whereas we could not detect any effect of PPP on clearance. The differing effects of exploitation and PPP imply that there can be different means by which variation in virulence is related to clearance, which could critically affect pathogen transmission and the evolution of pathogen virulence.

**Author summary:** Following an infection, hosts are not always able to quickly clear the pathogen, and they instead either die or survive with a persistent infection. Such variation is ecologically and evolutionarily important, because it can affect infection prevalence and transmission, and also virulence evolution. But what causes variation in infection outcomes? Here we contribute towards answering this question by investigating infection dynamics in flies infected with one of four bacterial species. We first establish that the bacterial species differ in virulence, *i.e*., the host death rate after infection. We find that variation in virulence arises because the bacteria differ in the two components of virulence: bacterial growth inside the host (exploitation), and the amount of damage caused per bacterium (per parasite pathogenicity).

Furthermore, as early-phase exploitation increases, bacterial clearance later in the infection decreases. This finding can be explained by increasing exploitation leading to increasing clearance costs for the host. Taken together we demonstrate that variation in infection outcomes can be partly explained by how virulence, and its components, relate to the rate of pathogen clearance. We propose that the decomposition of virulence is valuable for understanding variation in infection outcomes – potentially also beyond the interrelation between virulence and clearance.

## 1. Introduction

Once a host has become infected, the immune system will potentially limit pathogen growth, a response termed host resistance [1–3]. Resistance can therefore be quantified as the inverse of pathogen load [2]. Although there are clear benefits to the host of being able to mount an immune response that suppresses pathogen growth, resistance can come with evolutionary [4, 5] and usage costs [6] for the host [reviewed in 7]. During infection, hosts may re-allocate resources from other life history traits, such as reproduction [8] or development [9], into mounting an immune response. Furthermore, immune responses can lead to self-inflicted damage to the host, namely immunopathology [10–12]. Therefore, whether a pathogen is eliminated or not, *i.e.,* persists, is likely to depend upon the costs of infection *versus* the costs and effectiveness of the immune response against the infection, in addition to how well the pathogen can survive and replicate in the host environment.

Across host taxa, there is ample evidence of persistent bacterial infections. For example bacterial infections caused by *Escherichia coli* and *Staphylococcus aureus* can evade the human immune system and persist inside the host [13]. After injection with bacteria, insects can also sustain persistent systemic infections, for example in the mosquito *Anopheles gambiae* [14], the fruit fly *D. melanogaster* [15, 16] and the yellow mealworm beetle *Tenebrio molitor* [17]. These experimentally-induced infections can persist for at least 28 days in both *T. molitor* [17] and *D. melanogaster* [18], although longer term estimates are lacking.

Disparate bacterial species have been shown to be able to chronically infect (here defined as a minimum of seven days) the host species used in this study, *D. melanogaster* [15, 16, 18–23]. Persistent infections could be influential because the inability to clear an infection will result in more infected individuals in a population, thereby potentially increasing the probability of pathogen transmission. In the chronic infection phase in *D. melanogaster*, the bacterial load has been shown to stabilise around a relatively constant pathogen load over time [16, 21], which has been termed the set point bacterial load [SPBL; 21], after the set point viral load [*e.g*., 24]. However a stable infection load over time is not necessarily always the case, as the load for some bacterial species can gradually reduce in the days following infection, for example, in *D. melanogaster* injected with *E. coli* [20] and *T. molitor* injected with *S. aureus* [17, 25]. Alternatively, after an initial decline the infection load can start to increase again, as seen in the burying beetle, *Nicrophorus vespilloides*, injected with *Photorhabdus luminescens* [26].

Bacterial clearance during the chronic infection phase has not been commonly reported in insects and it may be related to the costs and benefits of immune system activation, and the degree of pathogen virulence. Virulence can be defined as disease severity, given as the decrease in host fitness caused by a pathogen [27], and which we here measure as reduced host survival. Virulence will be influenced by both host and parasite traits, *i.e*., it depends on resistance and tolerance from the host side, and the ability of the parasite to replicate and cause damage to the host [28]. From the pathogen perspective, variation in virulence across parasite strains could be due to differences in exploitation, that is an increase in virulence is a side effect of an increase in pathogen load [28, 29] (Fig 1A). However, variation in virulence could also be due to differences in per-parasite pathogenicity (PPP), *i.e*., the damage inflicted by each individual pathogen. PPP can be quantified by the slope of the reaction norm linking infection intensity (pathogen load) and host fitness [28, 29] (Fig 1B). A parasite genotype causing a steeper negative slope across a range of infection intensities, suggests higher PPP compared to a parasite genotype infection resulting in a shallower slope (Fig 1B). Here we use the concepts of exploitation and PPP to disentangle the causes of variation in virulence caused by infection with different bacterial species. We then use this information to uncover whether exploitation and PPP link to bacterial clearance probability.

**Fig 1.**
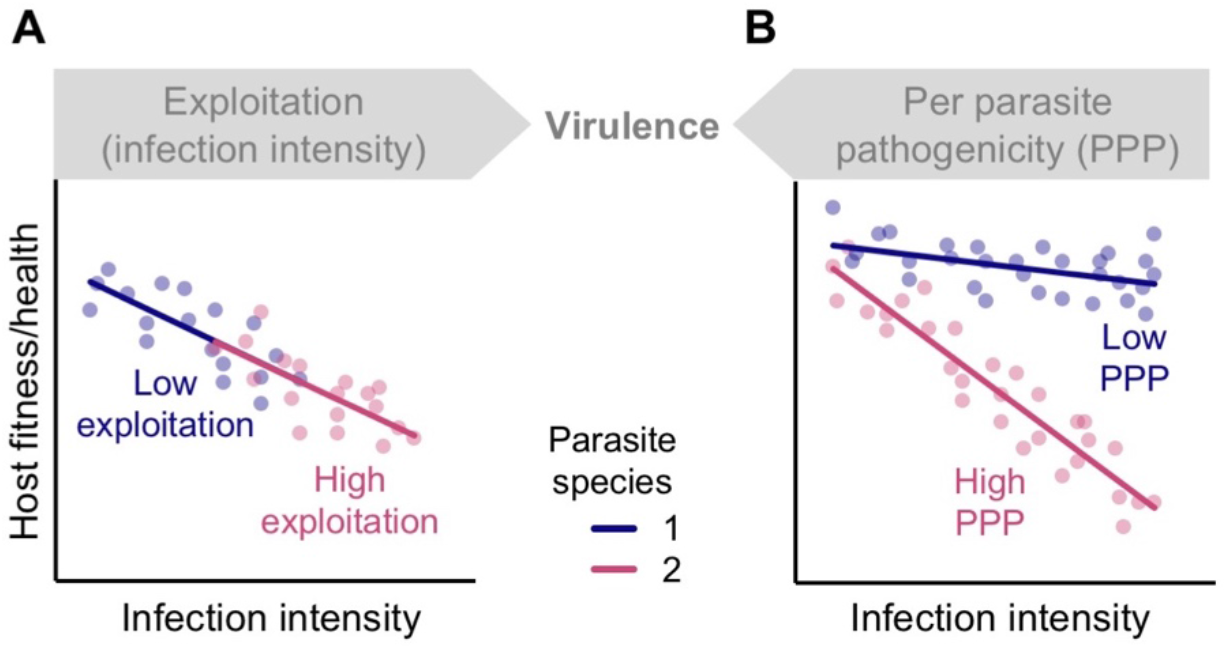
Decomposing virulence. Both exploitation and per parasite pathogenicity (PPP) can harm the host and thereby contribute towards virulence. Exploitation describes the infection intensity or bacterial load inside the host. PPP describes the damage per parasite that an infection does to the host. In this figure we show variation among pathogen species in exploitation and PPP, as illustrated by hypothetical relationships between host fitness and infection intensity for two species of parasite infecting the same host genetic background. **A.** In this example, the two parasite species have the same PPP but differ in exploitation. Parasite 1 has lower exploitation compared with parasite 2, because it causes a lower infection intensity. **B.** In contrast with A, in this example, parasite species have the same average exploitation but species 1 has lower PPP because its reaction norm has a shallower slope. This means that compared with species 2, species 1 causes less damage to the host with increasing parasite load. Figure modified from [29].

Exploitation and PPP are conceptually and mechanistically distinct ways in which parasites may harm their hosts. Accordingly, there could be distinct evolutionary consequences – both for pathogen and host traits. In the pathogen, reduced PPP and thus reduced virulence may be selected for, because this would result in a longer host infectious period without lowering the transmission rate [28]. On the other hand, higher exploitation might be selected for given that higher infection intensities are predicted to increase transmission rates and to take longer to clear (described as the recovery rate). However, increased exploitation and longer clearance will also mean increased virulence and hence trade-offs are expected between virulence, transmission and clearance [28, 30].

We hypothesise that exploitation and PPP can affect clearance in different ways, which could result in different, and potentially even opposing, patterns of how variation in virulence is related to clearance. In the following we consider how hosts are expected to react to infections with different levels of exploitation, or different levels of PPP. When the costs of mounting an immune response exceed the benefits of clearing an infection, one might predict a host to manage a persistent infection [31], and vice versa for clearing an infection. Therefore, we assume that hosts are more likely to clear an infection when the ratio of benefits to costs of clearance increases. In addition, as will be explained in more detail below, we assume that exploitation and PPP have different effects on the costs and benefits of clearance.

We assume that PPP mainly affects the benefits of clearance. Higher PPP directly results in a higher host death rate, which makes it more beneficial for the host to clear an infection. Accordingly, we expect that an increase in PPP leads to an increased effort by the host to clear the infection, which should result in a higher clearance rate (S1A Fig). In this context it is important to consider that an increased clearance effort could also lead to increased immunopathological effects, *i.e*., increased clearance costs in the form of increased virulence. Accordingly, such increased virulence would result in an increase in PPP (S1A Fig). This would mean that the measured PPP could reflect both the pathogen contribution to virulence and also the resulting host contribution to virulence. However, this potentially circular effect does not change our expectation for how PPP should be qualitatively related to clearance. Accordingly, we predict that an increase in measured PPP will be related to an increase in measured clearance.

In contrast to PPP, we assume that exploitation, *i.e*., infection intensity, affects both the benefits and also the costs of clearance. Increasing exploitation results in increased virulence, which increases clearance benefits. However, increasing exploitation additionally increases the costs of clearance if we assume that it is more difficult, and thus costlier, to clear infections with higher pathogen loads. Due to this dual increase in benefits and costs, two opposing predictions can be derived. First, if benefits of clearance increase faster than costs, we predict that increasing exploitation results in increasing clearance (S1B Fig). This prediction is consistent with observations of viral infections in humans and non-human primates where larger viral loads led to a faster decline in viral load or in shorter durations of viremia [reviewed in 32; e.g., 33]. Second, if costs of clearance increase faster than benefits, then we predict that increasing exploitation results in decreasing clearance (S1B Fig).

The initial exposure dose will also determine the outcome of infection, partly because microbe density at the beginning of an infection can determine the strength of the immune response [34]. Not only is dose-dependent survival frequently reported in response to bacterial infections [19, 26, 35], but bacterial load later in the infection has been demonstrated to correlate with the initial injection dose [19, 21]. We here test the generality of the latter finding. Furthermore, we predict that lower injection doses are more likely to be cleared, because a smaller population would be more susceptible to being killed by the host immune defences. The latter has been referred to as the inoculum effect in the context of *in vitro* antimicrobial activity studies [36–38]. However, the extent to which this kind of pattern is generalisable to bacterial infections *in vivo* is poorly understood.

Here we injected flies with four candidate bacterial species at a range of infection doses to address a number of different aims. First, we documented the variation in virulence and clearance caused by different bacterial species. Specifically, we tested whether species varied in virulence, which was measured via survival. We then decomposed virulence into its two constituent parts, to ask whether the species-level differences in bacterial virulence that we observed, were due to variation in parasite exploitation (infection intensity) or due to variation in PPP. To investigate clearance, we tested whether all four bacterial species establish a persistent infection by assessing infection status up to 35 days post injection, which is longer than any studies that we are aware of. In the second part we then tested our predictions regarding how variation in clearance is related to (1) exploitation and PPP, and (2) infection dose. Lastly, we assessed the infection status of flies that had died up to ten weeks post injection, which allowed us to assess whether long-term persistent bacterial infections occur in insects.

## 2. Materials and Methods

### 2.1. Fly population and maintenance

We used an outbred population of *Drosophila melanogaster* established from 160 *Wolbachia*-infected fertilised females collected in Azeitão, Portugal [39], and given to us by Élio Sucena. For at least 13 generations prior to the start of the experiments the flies were maintained on standard sugar yeast agar medium [SYA medium: 970 ml water, 100 g brewer’s yeast, 50 g sugar, 15 g agar, 30 ml 10 % Nipagin solution and 3 ml propionic acid; 40], in a population cage containing at least 5,000 flies, with non-overlapping generations of 15 days. They were maintained at 24.3 ± 0.2°C, on a 12:12 hours light-dark cycle, at 60-80 % relative humidity. The experimental flies were kept under the same conditions.

### 2.2. Bacterial species

We used the Gram-positive *Lactococcus lactis* (gift from Brian Lazzaro), Gram negative *Enterobacter cloacae subsp. dissolvens* (hereafter called *E. cloacae*; German collection of microorganisms and cell cultures, DSMZ; type strain: DSM-16657), *Providencia burhodogranariea* strain B (gift from Brian Lazzaro, DSMZ; type strain: DSM-19968) and *Pseudomonas entomophila* (gift from Bruno Lemaitre). *L. lactis* [41], *Pr. Burhodogranariea* [42] and *Ps. entomophila* [43] were isolated from wild-collected *D. melanogaster* and can be considered as opportunistic pathogens. *E. cloacae* was isolated from a maize plant, but has been detected in the microbiota of *D. melanogaster* [44]. These bacterial species were chosen based on various studies, which together suggest that they may be expected to show a range of virulence [18, 20, 21, 45–47].

### 2.3. Experimental design

For each bacterial species, flies were exposed to one of seven treatments: no injection (naïve), injection with *Drosophila* Ringer’s (injection control) or injection with one of five concentrations of bacteria ranging from 5 x 10^6^ to 5 x 10^9^ colony forming units (CFUs)/mL, corresponding to doses of approximately 92, 920, 1,840, 9200 and 92,000 CFUs per fly. The injections were done in a randomised block design by two people. Each bacterial species was tested in three independent experimental replicates. Per experimental replicate we treated 252 flies, giving a total of 756 flies per bacterium (including naïve and Ringer’s injection control flies). Per experimental replicate and treatment, 36 flies were checked daily for survival until all of the flies were dead. A sub-set of the dead flies were homogenised upon death to test whether the infection had been cleared before death or not. To evaluate bacterial load in living flies, per experimental replicate, four of the flies were homogenised per treatment, for each of nine time points: one, two, three, four, seven, 14, 21, 28- and 35-days post-injection.

### 2.4. Infection assay

Bacterial preparation was performed as in Kutzer *et al.* [18], except that we grew two overnight liquid cultures of bacteria per species, which were incubated overnight for approximately 15 hours at 30 °C and 200 rpm. The overnight cultures were centrifuged at 2880 rcf at 4 °C for 10 minutes and the supernatant removed. The bacteria were washed twice in 45 mL sterile *Drosophila* Ringer’s solution [182 mmol·L-1 KCl; 46 mol·L-1 NaCl; 3 mmol·L-1 CaCl2; 10 mmol·L-1 Tris·HCl; 48] by centrifugation at 2880 rcf at 4°C for 10 minutes. The cultures from the two flasks were combined into a single bacterial solution and the optical density (OD) of 500 µL of the solution was measured in a Ultrospec 10 classic (Amersham) at 600 nm. The concentration of the solution was adjusted to that required for each injection dose, based on preliminary experiments where a range of ODs between 0.1 and 0.7 were serially diluted and plated to estimate the number of CFUs. Additionally, to confirm *post hoc* the concentration estimated by the OD, we serially diluted to 1:10^7^ and plated the bacterial solution three times and counted the number of CFUs.

The experimental flies were reared at constant larval density for one generation prior to the start of the experiments. Grape juice agar plates (50 g agar, 600 mL red grape juice, 42 mL Nipagin [10 % w/v solution] and 1.1 L water) were smeared with a thin layer of active yeast paste and placed inside the population cage for egg laying and removed 24 hours later. The plates were incubated overnight then first instar larvae were collected and placed into plastic vials (95 x 25 mm) containing 7 ml of SYA medium. Each vial contained 100 larvae to maintain a constant density during development. One day after the start of adult eclosion, the flies were placed in fresh food vials in groups of five males and five females, after four days the females were randomly allocated to treatment groups.

Before injection, females were anesthetised with CO_2_ for a maximum of five minutes and injected in the lateral side of the thorax using a fine glass capillary (Ø 0.5 mm, Drummond), pulled to a fine tip with a Narishige PC-10, and then connected to a Nanoject II™ injector (Drummond). A volume of 18.4 nL of bacterial solution, or *Drosophila* Ringer’s solution as a control, was injected into each fly. Full controls, *i.e.*, naïve flies, underwent the same procedure but without any injection. After being treated, flies were placed in groups of six into new vials containing SYA medium, and then transferred into new vials every 2-5 days.

At the end of each experimental replicate, 50 µL of the aliquots of bacteria that had been used for injections were plated on LB agar to check for potential contamination. No bacteria grew from the Ringer’s solution and there was no evidence of contamination in any of the bacterial replicates. In addition, to confirm the concentration of the injected bacteria, serial dilutions were prepared and plated before and after the injections for each experimental replicate, and CFUs counted the following day.

### 2.5. Bacterial load of living flies

Flies were randomly allocated to the day at which they would be homogenised. Prior to homogenisation, the flies were briefly anesthetised with CO_2_ and removed from their vial. Each individual was placed in a 1.5 mL microcentrifuge tube containing 100 µL of pre-chilled LB media and one stainless steel bead (Ø 3 mm, Retsch) on ice. The microcentrifuge tubes were placed in a holder that had previously been chilled in the fridge at 4 °C for at least 30 minutes to reduce further growth of the bacteria. The holders were placed in a Retsch Mill (MM300) and the flies homogenised at a frequency of 20 Hz for 45 seconds. Then, the tubes were centrifuged at 420 rcf for one minute at 4 °C. After resuspending the solution, 80 µL of the homogenate from each fly was pipetted into a 96-well plate and then serially diluted 1:10 until 1:10^5^. Per fly, three droplets of 5 µL of every dilution were plated onto LB agar. Our lower detection limit with this method was around seven colony-forming units per fly. We consider bacterial clearance by the host to be when no CFUs were visible in any of the droplets. Although we note that clearance is indistinguishable from an infection that is below the detection limit. The plates were incubated at 28 °C and the numbers of CFUs were counted after ∼20 hours. Individual bacterial loads per fly were backcalculated using the average of the three droplets from the lowest countable dilution in the plate, which was usually between 10 and 60 CFUs per droplet.

*D. melanogaster* microbiota does not easily grow under the above culturing conditions [e.g., 47 and personal observation]. Nonetheless we homogenised control flies (Ringer’s injected and naïve) as a control. We rarely retrieved foreign CFUs after homogenising Ringer’s injected or naïve flies (23 out of 642 cases, *i.e*., 3.6 %). We also rarely observed contamination in the bacteria-injected flies: except for homogenates from 27 out of 1223 flies (2.2 %), colony morphology and colour were always consistent with the injected bacteria [see methods of 49]. Twenty one of these 27 flies were excluded from further analyses given that the contamination made counts of the injected bacteria unreliable; the remaining six flies had only one or two foreign CFUs in the most concentrated homogenate dilution, therefore these flies were included in further analyses. For *L. lactis* (70 out of 321 flies), *P. burhodogranaeria* (7 out of 381 flies) and *Ps. entomophila* (1 out of 71 flies) there were too many CFUs to count at the highest dilution. For these cases, we denoted the flies as having the highest countable number of CFUs found in any fly for that bacterium and at the highest dilution [20]. This will lead to an underestimate of the bacterial load in these flies.

### 2.6. Bacterial load of dead flies

For two periods of time in the chronic infection phase, *i.e*., between 14 and 35 days and 56 to 78 days post injection, dead flies were retrieved from their vial at the daily survival checks and homogenised in order to test whether they died whilst being infected, or whether they had cleared the infection before death. The fly homogenate was produced in the same way as for live flies, but we increased the dilution of the homogenate (1:1 to 1:10^12^) because we anticipated higher bacterial loads in the dead compared to the live flies. The higher dilution allowed us more easily to determine whether there was any obvious contamination from foreign CFUs or not. Because the flies may have died at any point in the 24 hours preceding the survival check, and the bacteria can potentially continue replicating after host death, we evaluated the infection status (yes/no) of dead flies instead of the number of CFUs. Dead flies were evaluated for two experimental replicates per bacteria, and 160 flies across the whole experiment. Similar to homogenisation of live flies, we rarely observed contamination from foreign CFUs in the homogenate of dead bacteria-injected flies (3 out of 160; 1.9 %); of these three flies, one fly had only one foreign CFU, so it was included in the analyses. Dead Ringer’s injected and naïve flies were also homogenised and plated as controls, with 6 out of 68 flies (8.8 %) resulting in the growth of unidentified CFUs.

### 2.7. Statistical analyses

Statistical analyses were performed in RStudio version 1.3.1073 [50]. The following packages were used for plotting the data: “ggpubr” [51], “grid”, “gridExtra” [52], “ggplot2” [53], “scales” [54], “survival” [55, 56] and “viridis” [57]. To include a factor as a random factor in a model it has been suggested that there should be more than five to six random-effect levels per random effect [58], so that there are sufficient levels to base an estimate of the variance of the population of effects [59]. In our experimental designs, the low numbers of levels within the factors ‘experimental replicate’ (two to three levels) and ‘person’ (two levels), meant that we therefore fitted them as fixed, rather than random factors [59]. However, for the analysis of clearance (see 2.7.7) we included species as a random effect because it was not possible to include it as a fixed effect due to the fact that PPP is already a species-level predictor.

#### 2.7.1. Do the bacterial species differ in virulence?

To test whether the bacterial species differed in virulence, we performed a linear model with the natural log of the maximum hazard as the dependent variable and bacterial species as a factor. Post-hoc multiple comparisons were performed using “emmeans”. The hazard function in survival analyses gives the instantaneous failure rate, and the maximum hazard gives the hazard at the point at which this rate is highest. We extracted maximum hazard values from time of death data for each bacterial species/dose/experimental replicate. Each maximum hazard per species/dose/experimental replicate was estimated from an average of 33 flies (a few flies were lost whilst being moved between vials etc.). To extract maximum hazard values we defined a function that used the “muhaz” package [60] to generate a smooth hazard function and then output the maximum hazard in a defined time window, as well as the time at which this maximum is reached. To assess the appropriate amount of smoothing, we tested and visualised results for four values (1, 2, 3 and 5) of the smoothing parameter, *b*, which was specified using *bw.grid* [61]. We present the results from *b* = 2, but all of the other values gave qualitatively similar results (see S2 Fig). We used *bw.method=“global”* to allow a constant smoothing parameter across all times. The defined time window was zero to 20 days post injection. We removed one replicate (92 CFU for *E. cloacae* infection) because there was no mortality in the first 20 days and therefore the maximum hazard could not be estimated. This gave final sizes of n = 14 for *E. cloacae* and n = 15 for each of the other three species.

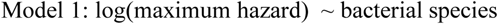

#### 2.7.2. Are virulence differences due to variation in parasite exploitation or PPP?

To test whether the bacterial species vary in PPP, we performed a linear model with the natural log of the maximum hazard as the dependent variable, bacterial species as a factor, and the natural log of infection intensity as a covariate. We also included the interaction between bacterial species and infection intensity: a significant interaction would indicate variation in the reaction norms, *i.e*., variation in PPP. The package “emmeans” [62] was used to test which of the reaction norms differed significantly from each other. We extracted maximum hazard values from time of death data for each bacterial species/dose/experimental replicate as described in 2.7.1. We also calculated the maximum hazard for the Ringer’s control groups, which gives the maximum hazard in the absence of infection (the y-intercept). We present the results from *b* = 2, but all of the other values gave qualitatively similar results (see results). We wanted to infer the causal effect of bacterial load upon host survival (and not the reverse), therefore we reasoned that the bacterial load measures should derive from flies homogenised before the maximum hazard had been reached. For *E. cloacae*, *L. lactis*, and *Pr. burhodogranariea*, for all smoothing parameter values, the maximum hazard was reached after two days post injection, although for smoothing parameter value 1, there were four incidences where it was reached between 1.8- and 2-days post injection. Per species/dose/experimental replicate we therefore calculated the geometric mean of infection intensity combined for days 1 and 2 post injection. In order to include flies with zero load, we added one to all load values before calculating the geometric mean. This was done using the R packages “dplyr” [63], “plyr” [64] and “psych” [65]. Each mean was calculated from the bacterial load of eight flies, except for four mean values for *E. cloacae*, which derived from four flies each.

For *Ps. entomophila* the maximum hazard was consistently reached at around day one post injection, meaning that bacterial sampling happened at around the time of the maximum hazard, and we therefore excluded this bacterial species from the analysis. We removed two replicates (Ringer’s and 92 CFU for *E. cloacae* infection) because there was no mortality in the first 20 days and therefore the maximum hazard could not be estimated. One replicate was removed because the maximum hazard occurred before day 1 for all *b* values (92,000 CFU for *E. cloacae*) and six replicates were removed because there were no bacterial load data available for day one (experimental replicate three of *L. lactis*). This gave final sample sizes of n = 15 for *E. cloacae* and n = 12 for *L. lactis*, and n = 18 for *Pr. burhodogranariea*.

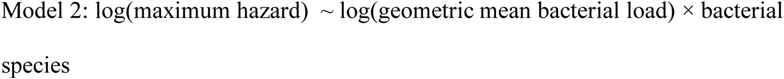

To test whether there is variation in parasite exploitation (infection intensity measured as bacterial load), we performed a linear model with the natural log of infection intensity as the dependent variable and bacterial species as a factor. Similar to the previous model, we used the geometric mean of infection intensity combined for days 1 and 2 post injection, for each bacterial species/dose/experimental replicate. The uninfected Ringer’s replicates were not included in this model. Post-hoc multiple comparisons were performed using “emmeans”. *Ps. entomophila* was excluded for the reason given above. The sample sizes per bacterial species were: n = 13 for *E. cloacae*, n = 10 for *L. lactis* and n = 15 for *Pr. burhodogranariea*.

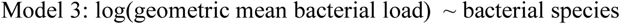

#### 2.7.3. Are persistent infection loads dose-dependent?

We tested whether initial injection dose is a predictor of bacterial load at seven days post injection [19, 21]. We removed all flies that had 0 CFU as they are not informative for this analysis. The response variable was natural log transformed bacterial load at seven days post-injection and the covariate was natural log transformed injection dose. Separate models were carried out for each bacterial species. Experimental replicate and person were fitted as fixed factors. By day seven none of the flies injected with 92,000 CFU of *L. lactis* were alive. The analysis was not possible for *Ps. entomophila* infected flies because all flies were dead by seven days post injection.

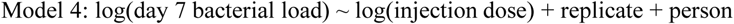

#### 2.7.4. Calculation of clearance indices

To facilitate the analyses of clearance we calculated clearance indices, which aggregate information about clearance into a single value for each bacterial species/dose/experimental replicate. All indices were based on the estimated proportion of cleared infections (defined as samples with measured zero bacterial load) of the whole initial population. For this purpose, we first used data on bacterial load in living flies to calculate the daily proportion of cleared infections in live flies for the days that we sampled. Then we used the data on fly survival to calculate the daily proportion of flies that were still alive. By multiplying the daily proportion of cleared flies in living flies with the proportion of flies that were still alive, we obtained the proportion of cleared infections of the whole initial population – for each day on which bacterial load was measured. We then used these data to calculate three different clearance indices, which we used for different analyses. For each index we calculated the mean clearance across several days. Specifically, the first index was calculated across days one and two post injection (clearance index_1,2_); the second index was calculated across days three and four (clearance index_3,4_); and the third index was calculated from days seven, 14 and 21 (clearance index_7,14,21_).

#### 2.7.5. Does injection dose affect bacterial clearance?

For this purpose, we aimed to assess clearance that occurs shortly after injection. Accordingly, we used the clearance index that was calculated for days one and two post injection. We suspected that the effect of dose on clearance might differ between bacterial species. Therefore, we ran separate tests for each species. The distribution of clearance values did not conform to the assumptions of linear models, therefore we used Spearman rank correlations to test whether clearance changes with injection dose.

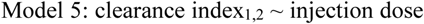

#### 2.7.6. Do the bacterial species differ in clearance?

To test whether the bacterial species differed in clearance, we used clearance index_3,4_, which is the latest timeframe for which we could calculate this index for all four species: due to the high virulence of *Ps. entomophila* we were not able to assess bacterial load and thus clearance for later days. The distribution of clearance values did not conform to the assumptions of a linear model. We therefore used a Kruskal-Wallis test with pairwise Mann-Whitney-U post hoc tests. To control for multiple testing we corrected the p-values of the post hoc tests using the method proposed by Benjamini and Hochberg [66] that is implemented in the R function *pairwise.wilcox.test*.

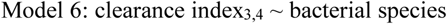

#### 2.7.7. Do exploitation or PPP predict variation in clearance?

To assess whether exploitation or PPP predict variation in clearance we performed separate analyses for clearance index_3,4_ and clearance index_7,14,21_. As discussed above, this precluded analysing *Ps. entomophila*. Exploitation and PPP were calculated based on bacterial load from days 1 and 2 (see section 2.7.2), therefore we did not perform an analysis for clearance index_1,2_. For each of the two indices we fitted a linear mixed effects model with the clearance index as the response variable. As fixed effects predictors we used the replicate-specific geometric mean log bacterial load (see section 2.7.2) and the species-specific PPP (see section 3.2). In addition, we included species as a random effect.

In our analysis we faced the challenge that many measured clearance values were at, or very close to zero. In addition, clearance values below zero do not make conceptual sense. To appropriately account for this issue, we used a logit link function (with Gaussian errors) in our model, which restricts the predicted clearance values to an interval between zero and one. Initial inspections of residuals indicated violations of the model assumption of homogenously distributed errors. To account for this problem, we included the log bacterial load and PPP as predictors of the error variance, which means that we used a model in which we relaxed the standard assumption of homogenous errors and account for heterogenous errors by fitting a function of how errors vary. For this purpose, we used the option *dispformula* when fitting the models with the function *glmmTMB*.

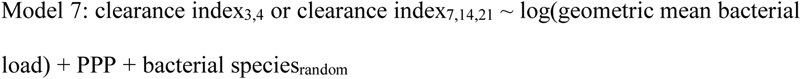

#### 2.7.8. Does longer-term clearance depend upon the injection dose?

In contrast to the analyses described above, we additionally aimed to assess the long-term dynamics of clearance based on the infection status of dead flies collected between 14 and 35 days and 56 to 78 days after injection. Using binomial logistic regressions, we tested whether initial injection dose affected the propensity for flies to clear an infection with *E. cloacae* or *Pr. burhodogranariea* before they died. The response variable was binary whereby 0 denoted that no CFUs grew from the homogenate and 1 denoted that CFUs did grow from the homogenate. Natural log transformed injection dose was included as a covariate as well as its interaction with day post injection, and person was fitted as a fixed factor. Replicate was included in the *Pr. burhodogranariea* analysis only, because of unequal sampling across replicates for *E. cloacae. L. lactis* injected flies were not analysed because only 4 out of 39 (10.3 %) cleared the infection. *Ps. entomophila* infected flies were not statistically analysed because of a low sample size (n = 12). The two bacterial species were analysed separately.

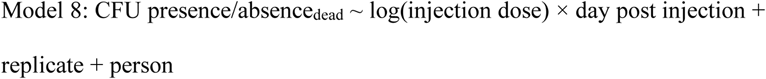

To test whether the patterns of clearance were similar for live and dead flies we tested whether the proportions of dead and live uninfected flies correlated with each other (S1 File).

## Results

### 3.1. Bacterial species vary in virulence

Fly survival after infection with five doses of the four different bacterial species is shown in Fig 2A-D. As predicted, the bacterial species differed significantly in virulence, given as the maximum hazard (F_3,55_ = 193.05, p < 0.0001; Fig 2E). All species differed significantly from each other (p < 0.0017 in all cases; S1 Table): *E. cloacae* was the least virulent and *Ps. entomophila* the most virulent bacterium. *Pr. burhodogranariea* and *L. lactis* were intermediate, with the former being less virulent than the latter. In all figures, the bacterial species are thus presented in order of virulence.

**Fig 2.**
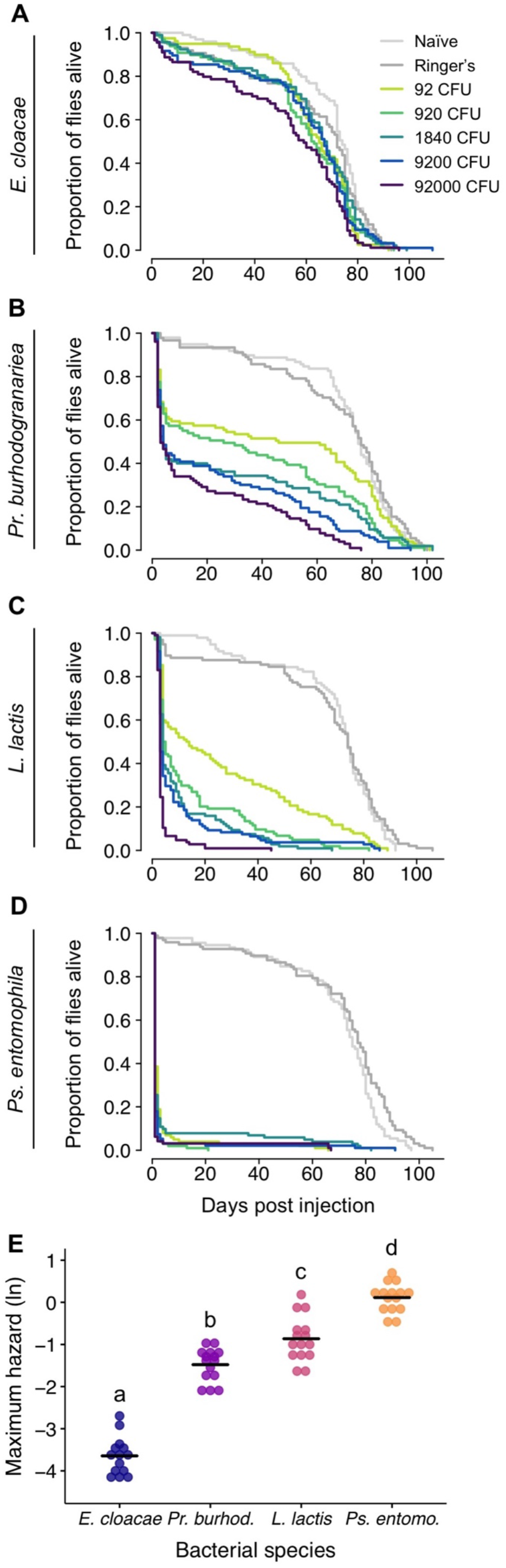
Fly survival after injection with one of four bacterial species. **A** – **D**. Survival curves after injection with four bacterial species, each at one of five doses. Controls were either injected with Ringer’s solution or received no injection (naïve). Each survival curve is from n = 79 to 108 flies. The legend in panel **A**, shows the treatments for all survival curves. **E**. The natural log of maximum hazard for all bacterial species, where each data point is the maximum hazard calculated from one replicate per dose. Ringer’s injected and naïve flies are not included. Black lines show means. Different letters denote means that are significantly different from one another (Tukey multiple comparison test).

### 3.2. Differences in virulence are due to variation in parasite exploitation and PPP

We assessed infection intensity over time post injection (Fig 3) and used the geometric mean of the values for the first two days post injection, as a proxy for parasite exploitation (see methods for rationale). Bacterial species varied significantly in exploitation of their hosts during this early infection phase (F_2,35_ = 35.90; p < 0.0001; Fig 4A). The least virulent bacterium, *E. cloacae*, had a significantly lower infection intensity, and thereby lower parasite exploitation, compared to either of the other species (Tukey contrasts: *E. cloacae vs. Pr. burhodogranariea*: t = −5.24, p < 0.0001; *E. cloacae vs. L. lactis*: t = −8.36, p < 0.0001). The more virulent bacterium, *L. lactis*, had the highest infection intensity, and differed significantly compared to the less virulent *Pr. burhodogranariea* (*L. lactis vs. Pr. burhodogranariea*: t = 3.50, p = 0.0018).

**Fig 3.**
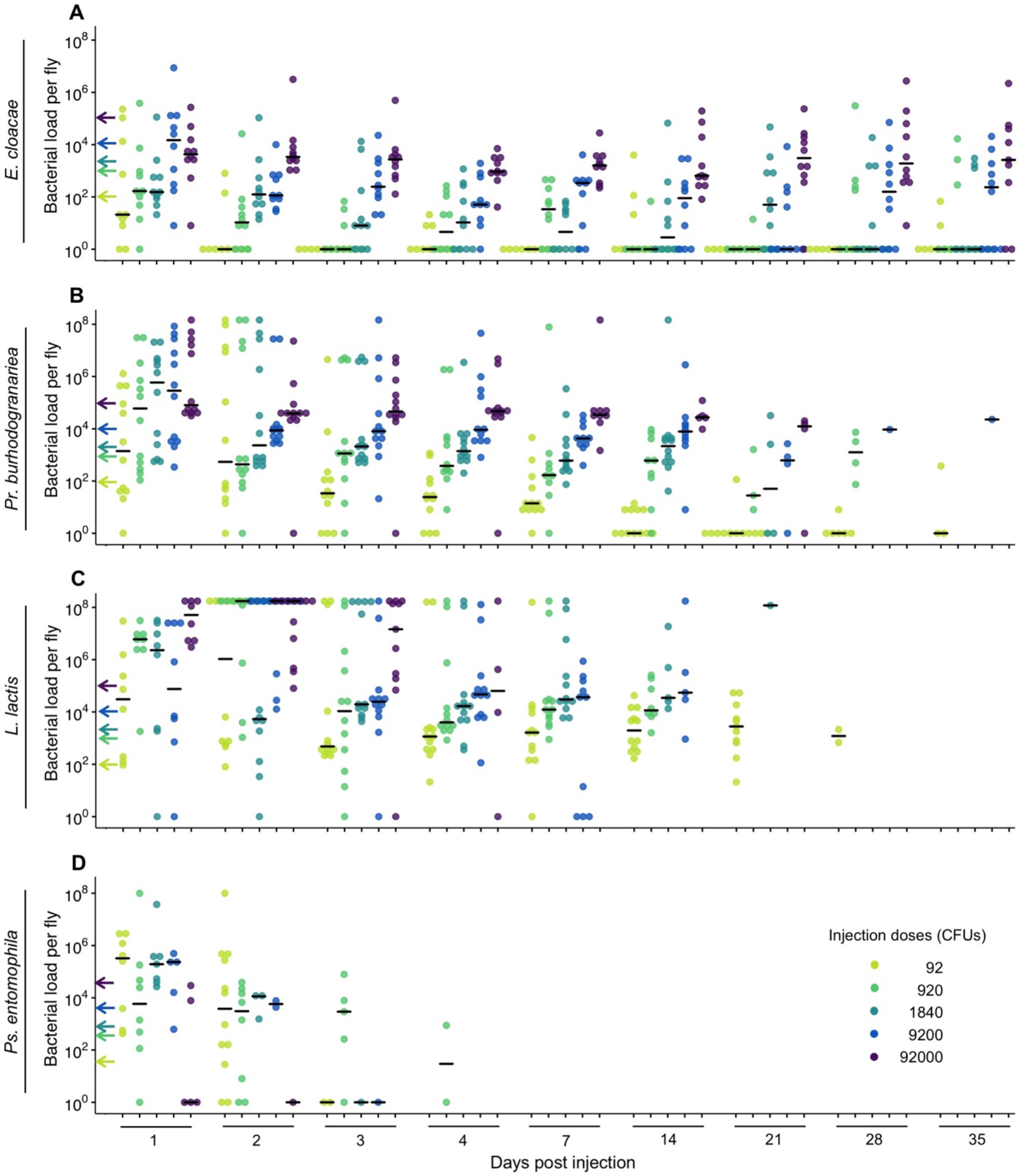
Bacterial load per living fly after injection with one of four bacterial species. (**A** – **D**). Flies were homogenised at between 1- and 35-days post-injection. The injection dose legend for all panels is shown in **D**. The arrows on the y-axis indicate the approximate injection doses. Missing data are due to increasing fly death over time. Black lines show medians.

**Fig 4.**
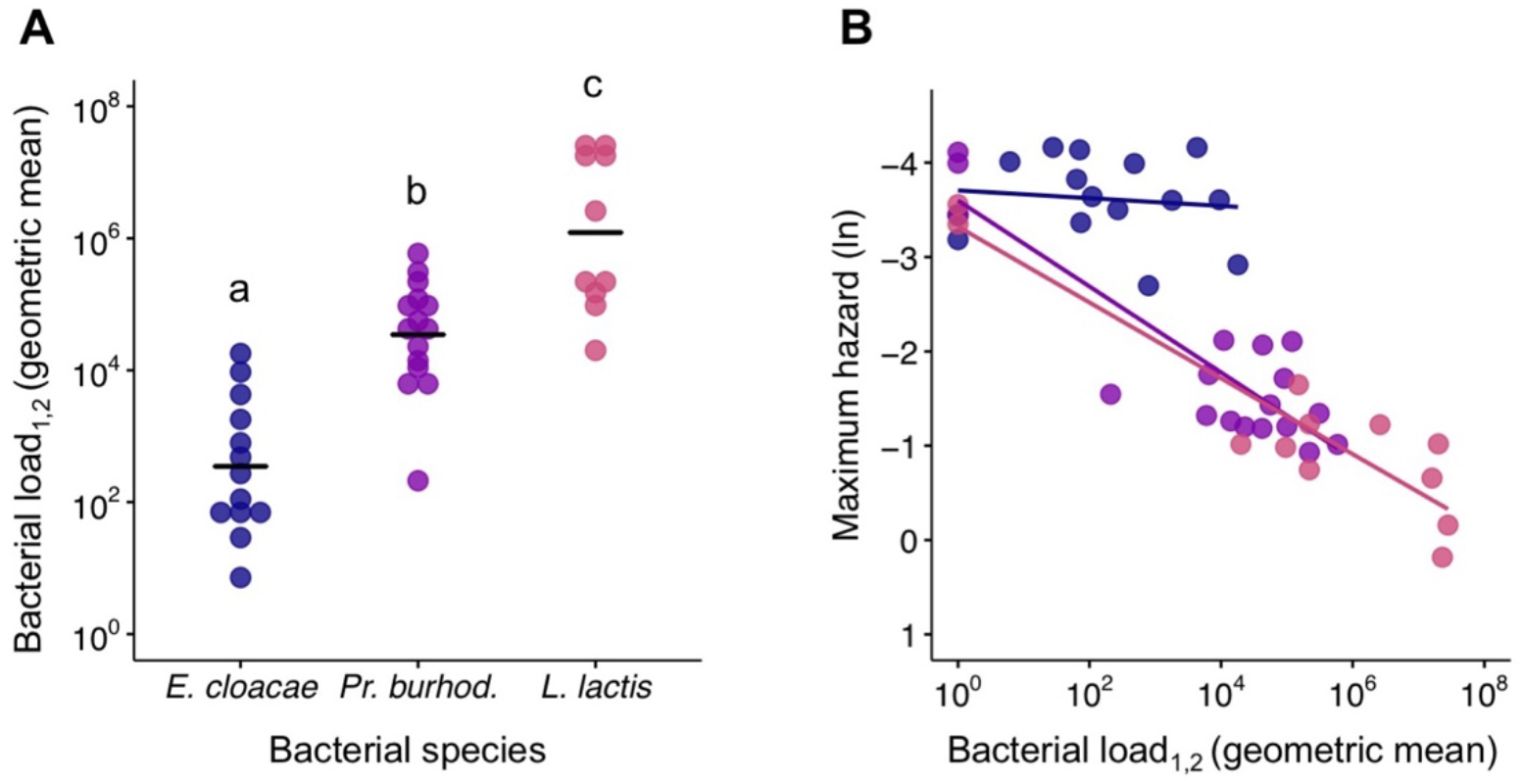
Virulence decomposition. **A**. Parasite exploitation given as infection intensity/bacteria load across bacterial species. Each data point is from one injection dose per bacteria, per experimental replicate, and gives the geometric mean of bacterial load for days one and two post injection (denoted as _1,2_). The circles are jittered along the x-axis to aid visualisation of overlapping data points. Black lines show means. Different letters denote means that are significantly different from one another (Tukey multiple comparison test). **B**. PPP given as the relationship between bacterial load and maximum hazard. The bacterial load data is the same as that given in **A** but with the addition of the Ringer’s control group. To allow inclusion of the uninfected Ringer’s control group to the figure we added one CFU to all mean bacterial load values. The natural log of maximum hazard data is estimated from survival data for the corresponding injection doses and experimental replicates. Maximum hazard is plotted as the inverse, such that the hazard (virulence) increases with proximity to the x-axis. Lines show linear regressions.

The slopes of the relationship between infection intensity and maximum hazard differed significantly across bacterial species, suggesting that the bacterial species differed in their PPP (infection intensity × bacterial species: F_2,39_ = 7.35, p = 0.0020; Fig 4B). *E. cloacae* had a relatively flat reaction norm, indicating a minimal increase in hazard with an increase in bacterial load, and thus a significantly lower PPP compared to both *Pr. burhodogranariea* (Tukey contrast: t = −3.74; p = 0.0017) and *L. lactis* (t = −3.34; p = 0.0052). In contrast, the latter two species had similar PPP to each other (t = −0.68; p = 0.78); both species had negative reaction norms, indicating an increase in hazard with an increase in bacterial load. There was no significant effect of bacterial load (F_1,39_ = 0.19, p = 0.67) or bacterial species F_2,39_ = 0.50, p = 0.61) on the maximum hazard. Qualitatively similar results were obtained using the three alternative smoothing parameters (S2 Fig). *Ps. entomophila* was not included in the above analyses because the maximum hazard was consistently reached at around day one post injection, meaning that we could not infer the causal effect of bacterial load upon survival.

### 3.3. All bacterial species established persistent infections

By homogenising living flies, we found that the two bacterial species with lower virulence, *E. cloacae* and *Pr. burhodogranariea* were able to persist inside the fly until at least 35 days post injection (Figs 2A and 2B respectively). The persistence estimates for *L. lactis* (28 days; Fig 3C) and *Ps. entomophila* (four days; Fig 3D) were both shorter, because the high mortality caused by these bacterial species meant that we could not test later time points. However, by testing for the presence or absence of bacteria in homogenised dead flies, we found that infections could persist for considerably longer, *i.e.,* around two and a half months:

*E.* cloacae = 77 days, *Pr. burhodogranariea* = 78 days, *L. lactis* = 76 days and *Ps. entomophila* = 75 days (Fig 5A-D).

**Fig 5.**
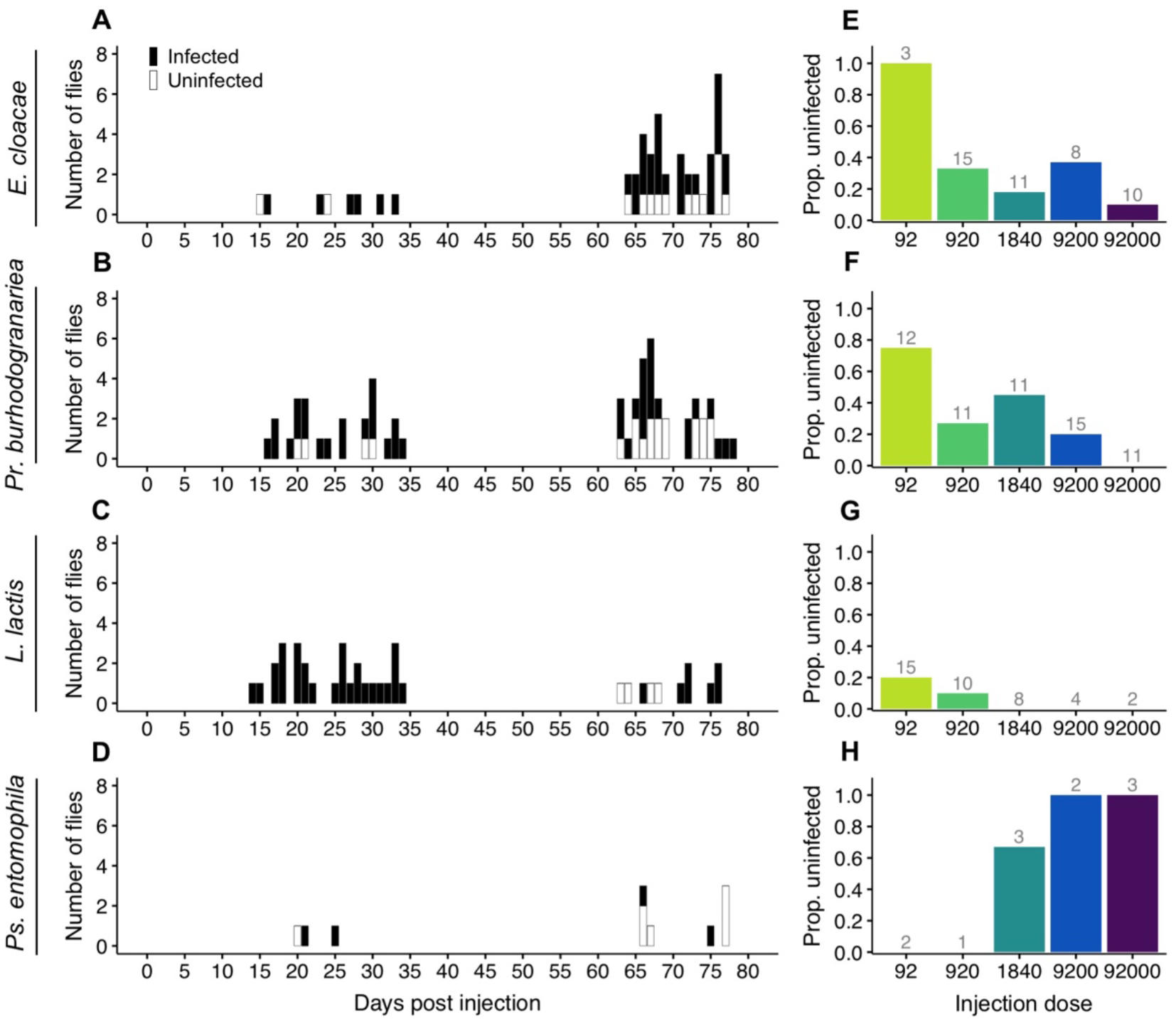
Bacterial clearance in dead flies. Each row shows flies that had been injected with one of four bacterial species. **A**-**D** The proportion of dead flies that were infected and uninfected according to the day post injection at which they died and were homogenised. Dead flies were homogenised at between 14 and 35, and 56 and 78, days post injection. **E**-**H** The same data as shown in the left-hand panels but graphed by injection dose. Numbers above the bars indicate the total numbers of flies from which the proportions were calculated, *i.e*., the total numbers of flies homogenised. Note that we cannot distinguish between flies that had cleared the infection and those where the bacterial load was below our detection limit.

### 3.4. Injection dose correlates with persistent infection loads

At seven days post injection, *E. cloacae* (Fig 6A) and *Pr. burhodogranariea* (Fig 6B) loads were significantly positively correlated with the initial injection dose (Table 1). *L. lactis* loads showed no significant correlation with the initial injection dose (Fig 6C; Table 1). We hypothesised that the lack of significant relationship between *L. lactis* load and injection dose might be due to our underestimation of the load of some flies injected with this bacterial species: this is because some flies had too many CFUs to count even at the lowest dilution, and they were therefore assigned a maximum bacterial load value (see methods), which was necessarily lower than their actual load. When we excluded the two flies at day seven that had been assigned the maximum value, the relationship became statistically significant (log(injection dose): F_1,35_ = 4.59; p = 0.039).

**Fig 6.**
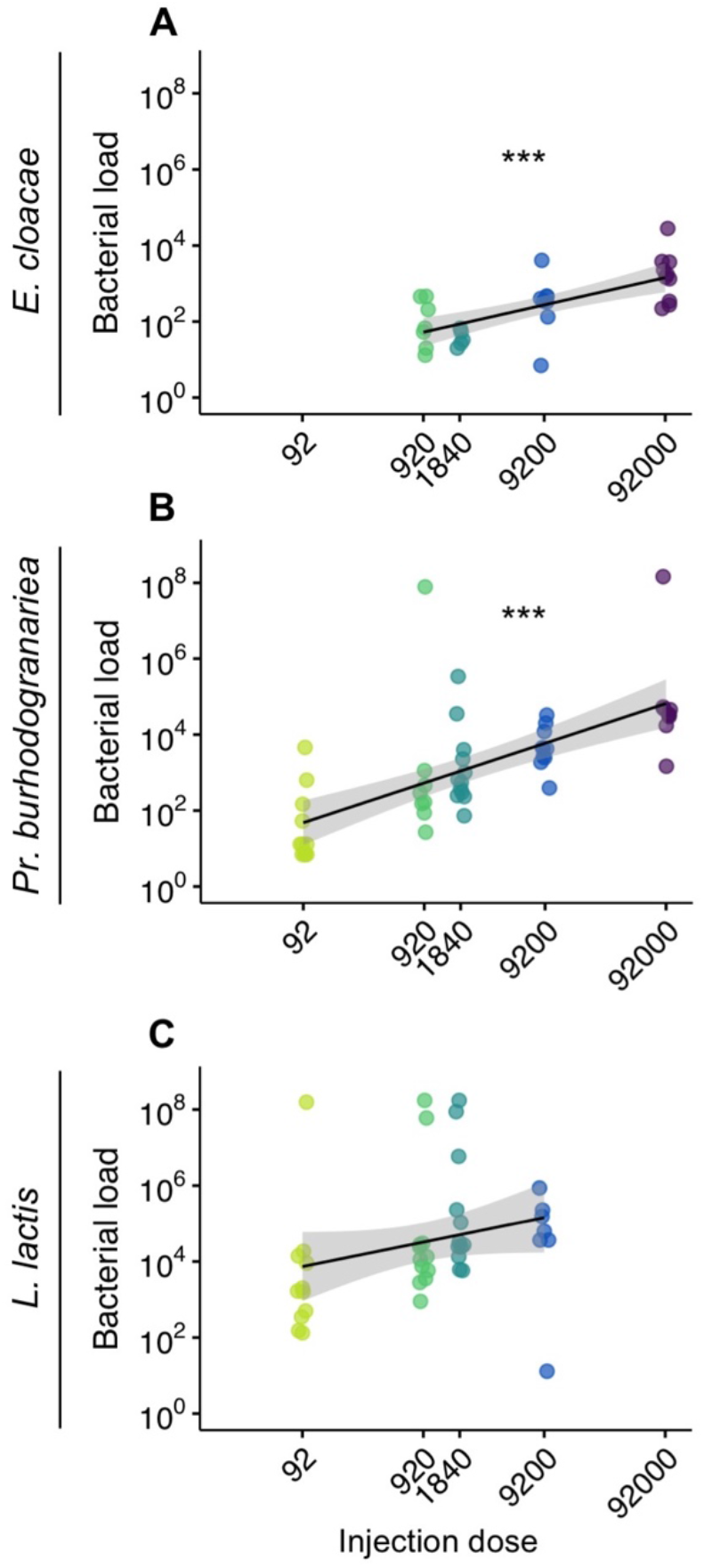
The relationship between bacterial load at seven days post injection, and the initial injection doses. Each row shows data from one bacterial species. Panel **A** contains no flies injected with 92 CFUs because all flies had a bacterial load of zero at day seven; **C** contains no flies injected with 92,000 CFUs because all flies had died by this time point. Each circle is the bacterial load of one fly, they are jittered along the x-axis to aid visualisation of overlapping data points, and they are coloured according to the injection dose. Flies with zero bacterial load are not shown (see methods). Linear regression lines are shown in black with 95 % confidence intervals. Asterisks denote significant correlations, where p < 0.0001.

**Table 1.** The effect of initial injection dose on bacterial load at seven days post injection (Model 4). Experimental replicate and the person performing the injection were also included as factors in the models. *Ps. entomophila* was not analysed because it caused high fly mortality. Statistically significant factors are in bold.

| Injected bacterium | Tested effect | df | F | P |
| --- | --- | --- | --- | --- |
| *E. cloacae* | Log(Injection dose) | 1,25 | 26.41 | **<0.0001** |
|  | Person | 1,25 | 0.16 | 0.69 |
|  | Replicate | 2,25 | 1.78 | 0.19 |
| *Pr. burhodogranariea* | Log(Injection dose) | 1,47 | 37.33 | **<0.0001** |
|  | Person | 1,47 | 0.23 | 0.63 |
|  | Replicate | 2,47 | 2.11 | 0.13 |
| *L. lactis* | Log(Injection dose) | 1,37 | 3.81 | 0.058 |
|  | Person | 1,37 | 0.71 | 0.40 |
|  | Replicate | 2,37 | 1.98 | 0.15 |

### 3.5. Lower doses of E. cloacae are cleared more quickly than higher doses

Summing up across all doses and days, 39.4 % (177 of 449) of *E. cloacae*-injected flies, 11.8 % (45 of 381) of *Pr. burhodogranariea*-injected flies, 3.7 % (11 of 301) of *L. lactis-*injected flies, and 21.4 % (15 of 70) of *Ps. entomophila*-injected flies cleared the infections (Fig 7). To account for mortality in our estimates of bacterial clearance we calculated clearance indices.

**Fig 7.**
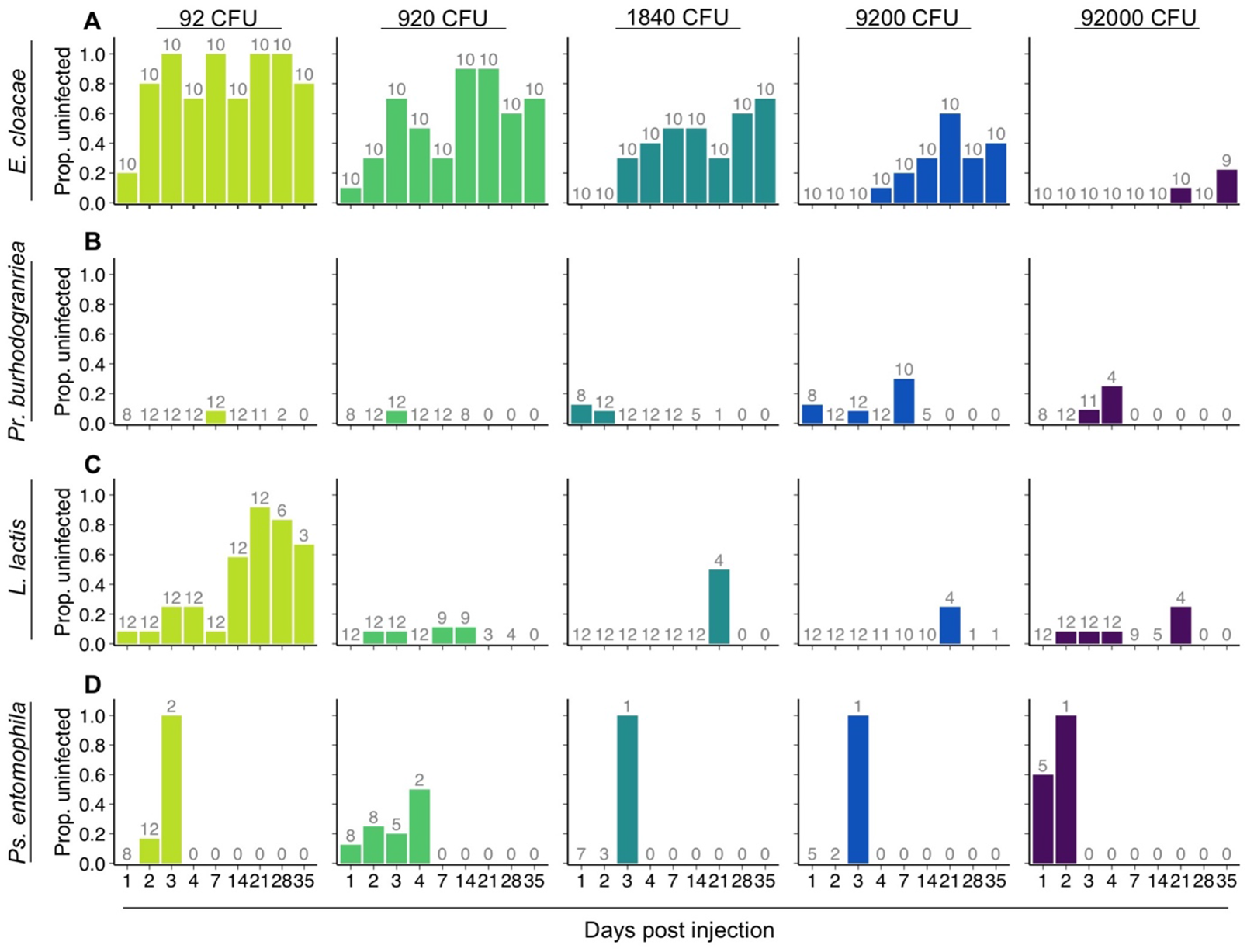
Bacterial clearance by living flies. Each row shows flies that had been injected with one of four bacterial species. **A**-**D** The proportion of live flies that were uninfected. Each column shows a different injection dose. Numbers above the bars indicate the total numbers of flies from which the proportions were calculated, *i.e.,* the total numbers of flies homogenised. Note that we cannot distinguish between flies that had cleared the infection and those where the bacterial load was below our detection limit.

Using the clearance index for the early infection phase, i.e., days one and two post injection (clearance index_1,2_), we found that flies injected with lower doses of *E. cloacae* were more likely to clear the infection compared to flies injected with higher doses (Spearman rank correlation: ρ = –0.86, p < 0.001, Fig 8A). However, the other three bacterial species did not show dose-dependent clearance (Fig 8B-D).

**Fig 8.**
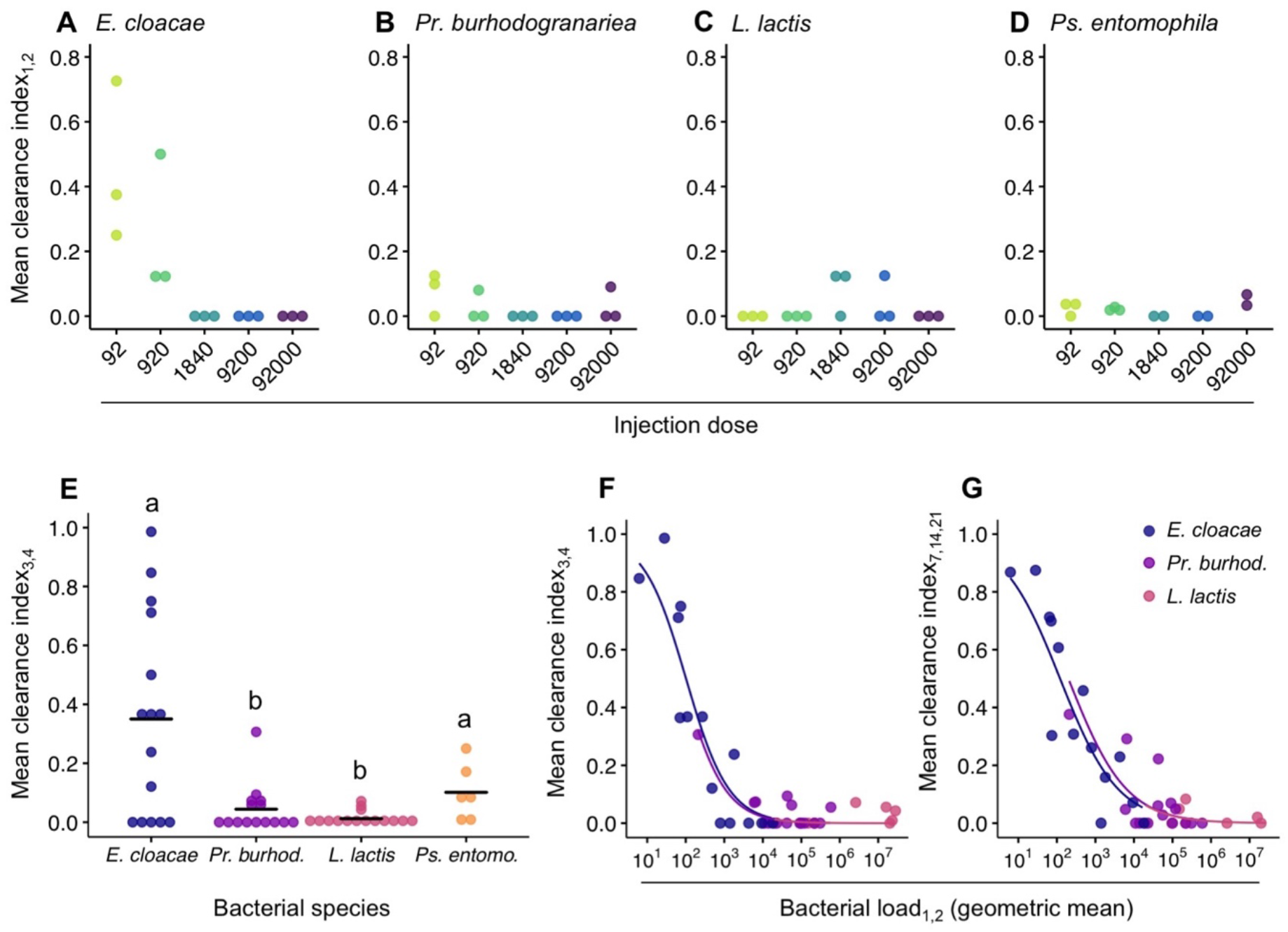
Effects of injection dose, species and exploitation on bacterial clearance. For all figures, each data point is from one injection dose per bacteria, per experimental replicate, and gives the mean proportion of cleared infections (out of the initial infected population) on days one and two (clearance index_1,2_), days three and four (clearance index_3,4_) or days seven, 14 and 21 (clearance index_7,14,21_). **A-D** Effect of injection dose on clearance index_1,2_ for each bacterial species. There was a statistically significant negative correlation for **A**. *E. cloacae* (ρ = –0.86, p < 0.001), but not for **B**. *Pr. burhodogranariea* (ρ = –0.36, p = 0.182), **C**. *L. lactis* (ρ = 0.13, p = 0.657) or **D**. *Ps. entomophila* (p = 0.982, ρ = –0.01). **E**. Mean species differences in clearance index_3,4_. The circles are jittered along the x-axis to aid visualisation of overlapping data points. Black lines show means. Different letters denote means that are significantly different from one another (Mann-Whitney-U post hoc tests). The effect of parasite exploitation, given as bacterial load, upon **F**. mean clearance index_3,4_ and **G**. mean clearance index_7,14,21_. The geometric mean of bacterial load was calculated from days 1 and 2 post injection (denoted as _1,2_), *i.e.*, the same values as in figure 3. There was a negative relationship between the two variables. Statistics are given in the main text and the species legend for both panels is shown in G.

### 3.6. Bacterial species vary in clearance

The four bacterial species used in this study covered a broad spectrum of clearance on days three and four post injection (Fig 8E). There was a statistically significant difference among species (p = 0.002, Chisq = 15.309, df = 3), and after p-value correction, the post hoc tests indicated statistically significant differences among the following species pairs: *E. cloacae* and *Pr. burhodogranariea* (p = 0.024), *E. cloacae* and *L. lactis* (p = 0.011), *Ps. entomophila* and *Pr. burhodogranariea* (p = 0.048), *Ps. entomophila* and *L. lactis* (p = 0.011).

Rather than matching the virulence gradient across species (Fig 2E), clearance formed a U-shaped pattern with the species with the highest virulence (*Ps. entomophila*) and lowest virulence (*E. cloacae*) showing higher levels of clearance compared to the two species of intermediate virulence (*Pr. burhodogranariea* and *L. lactis*).

### 3.7. Exploitation but not PPP predict clearance

Our analyses of clearance index_3,4_ and clearance index_7,14,21_ showed similar results. In both cases we found no statistically significant effect of PPP, but a significant negative effect of exploitation, such that as the mean bacterial load for days one and two increased, clearance decreased (Fig 8F, G Table 2). Similar results were obtained using the three alternative smoothing parameters for calculating PPP (S3 Fig).

**Table 2.** The effect of log bacterial load (exploitation) and PPP on two different clearance indices. Statistically significant factors are in bold.

| Response variable | Tested effect | df | Chisq | P |
| --- | --- | --- | --- | --- |
| **Clearance index<sub>3,4</sub>** | Log (geometric mean bacterial load) | 1 | 14.16 | < **0.001** |
|  | PPP | 1 | 0.38 | 0.535 |
| **Clearance index<sub>7,14,21</sub>** | Log (geometric mean bacterial load) | 1 | 36.34 | < **0.001** |
|  | PPP | 1 | 0.35 | 0.557 |

### 3.8. Bacterial clearance before death is dose dependent

We homogenised flies that died during the chronic phase of the infection (between 14 and 35 days and between 56- and 78-days post injection) to test whether they died whilst being infected, or whether they were able to clear the infection before death. Flies were indeed able to clear the infection before death, but the degree to which this occurred varied across bacterial species (Fig 5). Furthermore, for all bacterial species in both homogenisation phases there were flies where the infection persisted until death, and flies that were uninfected at death (Fig 5A-D). Lower injection doses of *E. cloacae* were more likely to be cleared before death than higher injection doses (Fig 5E; Table 3), but there was no significant effect for *Pr. burhodogranariea* (p = 0.051; Fig 5F; Table 3). Summing up across all doses and days, 29.8 % (14 out of 47) of *E. cloacae*-injected flies, 33.3 % (20 out of 60) of *Pr. burhodogranariea*-injected flies, 10.3 % (4 out of 39) of *L. lactis-*injected flies, and 66.7 % (8 out of 12) of *Ps. entomophila*-injected flies cleared the infection before death.

**Table 3.** The effect of injection dose on presence/absence of infection in dead flies (Model 8). Person performing the injection was also included as a factor in the models, and replicate was included for the analysis for *Pr. burhodogranariea* infections. Statistically significant factors are in bold.

| *Injected bacterium* | *Tested effect* | *df* | *LR Chisq* | *P* |
| --- | --- | --- | --- | --- |
| *E. cloacae* | Log(Injection dose) | 1 | 3.82 | 0.051 |
|  | Day post injection | 1 | 0.0037 | 0.95 |
|  | Person | 1 | 1.22 | 0.27 |
|  | Log(Injection dose) ×<br>Day post injection | 1 | 0.013 | 0.91 |
| *Pr. burhodogranariea* | Log(Injection dose) | 1 | 13.26 | **0.00027** |
|  | Day post injection | 1 | 2.99 | 0.084 |
|  | Person | 1 | 4.33 | **0.037** |
|  | Replicate | 1 | 0.038 | 0.54 |
|  | Log(Injection dose) ×<br>Day post injection | 1 | 1.75 | 0.19 |

## Discussion

In this study we demonstrate that sustained persistent infection and clearance are both possible outcomes for bacteria showing a range of virulence when they infect female *D. melanogaster*. We show that bacteria of all species can persist inside the host for at least 75 days and that bacterial virulence differences can be explained by a combination of variation in exploitation and PPP. We also show that lower doses of the bacterium with the lowest virulence, *E. cloacae*, are cleared more quickly than higher doses, and that clearance rates are species specific. Furthermore, higher exploitation of the host leads to lower bacterial clearance. Finally, we found some support for our hypothesis that exploitation and PPP imply different costs and benefits of clearance and thus both virulence components have different effects on clearance.

### 4.1. Differences in virulence are due to variation in exploitation and PPP

The infecting bacterial species showed pronounced differences in virulence. To understand why this was the case, we decomposed virulence into its two components: exploitation and PPP [28, 29]. Exploitation, given as infection intensity or bacterial load, is the more frequently tested explanation for variation in virulence [28]. There is ample evidence that exploitation varies across parasite genotypes [67, e.g., monarch butterflies and their protozoan parasites: 68, *Daphnia magna* infected with the bacterium *Pasteuria ramosa*: 69], and also, unsurprisingly, that it varies across parasite species infecting the same host genotype [19–21]. Indeed, in the current study, all bacterial species tested showed significant differences in exploitation, where bacterial load increased as virulence increased. Chambers et al. [19] observed that the two bacterial species in their study that caused lower mortality showed little initial proliferation inside the host, but that the species causing more mortality showed an initial increase in the bacterial load: these results support our findings.

However, variation in virulence might not only be determined by the load that a pathogen attains: Råberg & Stjernman [28] proposed that pathogen genotypes may also vary in PPP *i.e*., the harm or damage caused per parasite [28, 29]. Some pathogens may cause more damage to the host independently of their density, through mechanisms that directly affect the host homeostasis. For example, *Bacillus anthracis* produces two anthrax toxins which are responsible for impairing the host immune system and disrupting basic cellular functions, ultimately killing the host [70]. By comparison, the genetically similar *Bacillus cereus* usually only produces a mildly virulent gastro-intestinal infection [71–73]. Variation in PPP can be observed when different parasite genotypes show different reaction norms for the relationship between host health and infection intensity, when infecting the same host genotype [29]. Variation in PPP has been demonstrated for rats infected with different clones of *Plasmodium chabaudii,* the agent of rodent malaria [28], for different strains of protozoan parasites infecting monarch butterflies [67], and for humans infected with different HIV-1 genotypes [74]. Here we found a significant overall effect of PPP across bacterial species, whereby *Pr. burhodogranariea* and *L. lactis* had significantly more negative slopes compared to *E. cloacae*. This finding, combined with the exploitation results, implies that *E. cloacae* is less virulent towards its host compared to the other two species, because of a combination of lower PPP and less exploitation. On the other hand, given that *Pr. burhodogranariea* and *L. lactis* both showed similar levels of PPP, it suggests that the variation in virulence between these two species is due to higher exploitation by *L. lactis*, rather than differences in PPP. Some of our *L. lactis* counts were underestimates because there were too many CFUs to count, so it is possible that the slope of the reaction norm might have differed slightly had we not encountered this issue. Nonetheless, had we only examined exploitation as a source of variation, we might have concluded that load alone explains the differences that we found in virulence. These two sources of variation, exploitation and PPP, have not frequently been explored in the same study, so it is generally difficult to ascertain the relative importance of the two sources of variation. However, variation in PPP was demonstrated to explain more of the variance in virulence across HIV-1 genotypes than did set point viral load [74].

### 4.2. All bacterial species established persistent infections

All four bacterial species were able to establish persistent infections in *D. melanogaster*. *E. cloacae* and *Pr. burhodogranariea* could be retrieved from live homogenised flies up to 35 days, *L. lactis* up to 28 days, and *Ps. entomophila* up to four days post injection. The reduced estimates for the latter two species are due to higher mortality, meaning that no flies were alive to sample at later time points. However, by homogenising flies that had died, we show that all bacterial species can persist inside the host for at least 75 days. To the best of our knowledge these estimates are far beyond the currently known length of persistent infections after injection in insects [28 days: 17, 18]. The duration of infection can be of key ecological and potentially also evolutionary importance, because persistence determines the prevalence of infection in a population, and therefore could affect transmission.

It is unclear how these bacteria are able to persist for so long inside the host, although there are a number of theoretical possibilities, for example, through surviving inside host tissue, forming biofilms or existing as persister or tolerant cells [75]. *Salmonella typhimurium* [76] and *S. aureaus* [77] can survive inside insect haemocytes. The bacterial species that we used are not known to be intracellularly replicating, *e.g*., some *Providencia* strains were able to survive, but not replicate, at low numbers 24 hours after infecting a *D. melanogaster* S2 cell line [45]. Both *E. cloacae* and *L. lactis* are able to produce biofilms *in vitro* [78, 79, respectively], but it is unknown whether this is the case inside an insect host. *Pr. burhodogranariea* is not able to form biofilms *in vitro* [45] although it is unknown if it might still be possible *in vivo*. Biofilms can cause chronic infections such as *Pseudomonas aeruginosa* in cystic fibrosis patients [80], and oral infection of *D. melanogaster* with *Ps. aeruginosa* resulted in biofilm production in the crop [81], but it is unknown whether *Ps. entomophila* forms biofilms *in vivo*. Lastly, bacteria could potentially survive inside the host in persistent or tolerant cell states, as is discussed in relation to the failure of antibiotic treatments [82]. However, persistent cells typically make up a small proportion (< 1%) of the bacterial population [82], so this might not explain the high numbers of CFUs that are retrieved from the flies. Future research will test the likelihood of these and other mechanisms.

*D. melanogaster* that are able to control a bacterial infection during the acute infection phase have been shown to have a relatively constant bacterial load in the chronic infection phase, which Duneau *et al.* [21] found remains stable until at least ten days post injection for *Pr. rettgeri*. Our bacterial load data (Fig 3) suggests that *E. cloacae*, *Pr. burhodogranariea* and *L. lactis*, show relatively stable loads from around day three to four post infection, lending support to the SPBL concept. In addition, Duneau *et al*. [21] observed that per host, bacteria with low virulence had a SPBL of a few hundred bacteria, whereas bacteria of intermediate virulence had a SPBL of a few thousand bacteria. Our data also lend support to the idea that virulence relates to SPBL, given that low virulence *E. cloacae* had a persistent load of tens to hundreds of bacteria, and high virulence *L. lactis* had a load of tens of thousands of bacteria. This finding is supported by the virulence decomposition analysis (see section 4.1), which shows that as virulence increases, so does exploitation of the host over the first couple of days post infection. Therefore, more virulent bacteria have higher initial proliferation rates as shown by exploitation. Given that the infection load stays relatively constant in the longer term, the initial proliferation differences likely explain the relationship between SPBL and virulence.

### 4.3. Injection dose correlates with persistent infection loads

The bacterial load at day seven post injection, positively correlated with the initial injection dose for *E. cloacae, Pr. burhodogranariea* and *L. lactis* (but see results section for the latter). Our results expand the known bacterial species for which this relationship exists, and they lend weight to the idea that this may be a more general phenomenon in *D. melanogaster* bacterial infections. Previous studies found that this relationship held for bacterial load at seven- and fourteen-days post injection [E. faecalis, Pr. rettgeri and S. marcescens: 19, Pr. rettgeri: 21;]. It has been suggested that the SPBL will remain at around the bacterial load at which the infection was controlled [19, 21]. Given that insects can show dose dependent inducible immune activation [34], and given that the antimicrobial peptide Drosocin has been shown to control *E. cloacae* infections and that a combination of Drosocin, Attacins and Diptericins control *Pr. burhodogranariea* infections [47], one could hypothesise that these AMPs are to some degree involved. However, the mechanisms that allow a dose-dependent persistent infection, remain to be uncovered. It was not possible to test *Ps. entomophila* given its high mortality during the acute infection phase.

### 4.4. Lower doses of E. cloacae are cleared more quickly than higher doses

The likelihood of clearing *E. cloacae* was dose dependent, although we did not find a dose threshold below which there was complete clearance in all flies. The finding of dose-dependent clearance, whilst maybe not surprising, could explain some discrepancies across studies in terms of whether evidence of persistent infections is found. Just as stochastic variation explains variation in the outcome of the early infection phase [21], perhaps stochasticity plays a role in the clearance of bacteria [83], particularly where infection loads are low such as in *E. cloacae*, for example through variation in expression of Drosocin. In comparison to *E. cloacae*, most replicates of the other species showed no clearance in the early infection phase, i.e., one- and two-days post infection, thus dose did not influence clearance for these species.

### 4.5. Bacterial species vary in clearance

Across species, we uncovered a U-shaped pattern in bacterial clearance, where the species with the lowest and highest virulence had higher levels of clearance compared to the two species of intermediate virulence. Our finding that the low virulent species could be cleared, is supported by evidence from infections with low virulence *E. coli* and *Erwinia carotovora Ecc15*, where an injection dose of 30,000 bacteria was cleared in 22 % and 8 % of flies, respectively [21]. The clearance of intermediate and high virulence pathogens in *D. melanogaster* has been described as being rare, because no bacteria were cleared from any of the previously infected hosts over the seven-days post injection [21]. *Pr. rettgeri* and *Enterococcus faecalis* were described as having intermediate virulence in that study, with a survival of around 50-60% seven days post infection [21], which is in the region of the survival that we found for our intermediate virulence species, *Pr. burhodogranariea* and *L. lactis*. Although clearance in our intermediately virulent bacteria was lower than that of the low and high virulent bacteria, our results challenge the finding that clearance is rare because our three more virulent bacteria all appear to be clearable to differing degrees, including within the first seven days of infection. Our results thereby show that persistent infections are not inevitable. This finding is supported by the observation that *Pr. burhodogranariea* was cleared in flies seven- to ten-days post injection [45; although low sample sizes]. Similarly to the current study, Kutzer & Armitage [20] also found that a few female flies, inoculated with a dose of *L. lactis* in common with this study (1,840 CFUs), cleared the infection (3 out of 141; 2.1 %). Lastly, we expected that there may be selection for a fast and efficient early clearance of infection by *Ps. entomophila*, because of its high virulence. Clearance of *Ps. entomophila* was indeed higher than for the intermediate bacteria, although mortality was too high to assess clearance in living flies for longer than four days post injection. Nonetheless, there is evidence from other studies that *Ps. entomophila* has high virulence and can be cleared from other *D. melanogaster* populations/genotypes [18, 39, 46]. The ability to resist *Ps. entomophila* infection varied across host genotypes; similarly, five out of the ten tested genotypes contained some individuals who cleared this pathogen [46]. Host genotypic variation in clearance may thereby more generally explain why some studies find clearance and others do not.

Even though dead flies were sampled for a longer period post-injection (up to 78 days) compared to live flies (up to 35 days), the patterns of bacterial clearance in dead flies largely reflected the results for live flies, given that dead flies that had been infected with *E. cloacae* showed dose dependency in clearance. Once again, comparatively few dead individuals had cleared *L. lactis* infections, whereas proportionally more had cleared *Ps. entomophila.* Because we processed dead flies up to 24 hours post-death we did not analyse the number of CFUs, however it would be interesting to test whether the bacterial load upon death [21] remains constant even after many weeks of a potentially costly infection.

### 4.6. Exploitation but not PPP predicts clearance

We next sought to understand how the two components that determine variation in virulence affect clearance of the infection, whilst framing our argumentation in the context of the benefits to costs ratio to the host of clearing an infection. Our results provide some support for our hypothesis that exploitation and PPP can affect clearance in different ways, which could result in different, and potentially even opposing, patterns of how variation in virulence is related to clearance. As predicted, we found that across species, increased host exploitation early in the infection, i.e., days one and two post infection, was associated with decreased clearance at days three and four post infection, and also at days seven to 21. This finding is consistent with the expectation that the host immune response is sensitive to clearance costs associated with exploitation. We did not find support for our prediction that increasing PPP would be associated with increased clearance, which would be expected if the host immune response is sensitive to clearance benefits. Taken together, our results demonstrate that for the investigated pathogens, variation in clearance is mainly explained by variation in clearance costs whereas clearance benefits seem to have no or only little effect. Further studies are necessary to assess whether this is a general pattern.

Our finding of a negative relationships between exploitation and clearance is in contrast to data from vertebrate viral infections [reviewed in 32;, e.g., 33], where larger viral loads led to a faster decline in viral load or in shorter durations of viremia. Although these studies did not directly assess clearance as defined in our study, their results are consistent with the idea that in these systems increasing exploitation might lead to a stronger increase in the benefits of clearance compared to the costs of clearance. In this case one would expect a positive relationship between exploitation and clearance.

In contrast to exploitation, by definition, PPP cannot vary within replicates of a species, because each species has just one PPP slope. In addition, only three species could be included in this analysis due to high mortality after injection with *Ps. entomophila*. Therefore, we note that the statistical power of detecting an effect of PPP on clearance might have been lower than the power to detect an effect of exploitation. Nonetheless, it is possible that analyses with more species will reveal an effect of PPP on clearance. Based on existing information on the biology of *Ps. entomophila* we propose that the increased levels of clearance for this species (Fig 8E) might have been caused by particularly high levels of PPP. *Ps. entomophila* produces the virulence factor AprA [84], and a pore-forming toxin called Monalysin [85] in association with the activation of stress-induced pathways and an increase in oxidative stress [86].

Ultimately this leads to a lack of tissue repair in the gut, and in most cases fly death [reviewed in 87]. If similar pathologies are induced in the haemocoel after infection, in contrast to other bacterial species, sustaining a persistent bacterial load in the face of high levels of tissue damage might rarely be a viable option. Instead, the fly host might activate a stronger immune response to attempt to clear the infection [31, 88]. However, a few flies in ours and other studies, did survive into the chronic phase whilst testing positive for a *Ps. entomophila* infection. In *D. melanogaster* Drosocrystallin expression in the gut has been shown to confer protection to Monalysin and lead to higher individual survival [89, 90]; perhaps individuals surviving into the chronic infection phase exhibit higher expression of such mechanisms, allowing them to tolerate the damage caused by infection. Nonetheless, we suggest that the higher virulence of *Ps. entomophila* could have been driven by an increased PPP because of the above-mentioned toxins, but additional experiments would be required to test this idea.

### 4.7. Conclusions

To conclude, our results strongly support that PPP is an important component driving variation in virulence, and that disentangling its contribution towards virulence, in combination with the contribution of exploitation, will undoubtedly help our mechanistic and evolutionary understanding of host-pathogen interactions. We suggest that such a decomposition of virulence can be used to better understand how virulence relates to other infection processes such as clearance during persistent infections. Future research will be needed to test the generality of the relationships we have uncovered between virulence decomposed as exploitation and PPP, and the persistence and clearance of infections. Here we have taken a pathogen perspective on virulence, however a more complete understanding of how virulence relates to clearance will be achieved by simultaneously considering the host factors that contribute to variation in virulence, i.e., resistance and tolerance. Such an approach may contribute towards expanding the theory on virulence evolution.

## Author contributions

SAOA conceived the overall idea. BAH, LS and SAOA designed the experiments and collected the data. BAH, LS, MF and SAOA wrote the manuscript. MF & RRR conceived the virulence decomposition and clearance analyses and MF, RRR & SAOA analysed the data. All authors contributed critically to the drafts.

## Acknowledgements

We thank the Hiesinger group, the Rolff group, Luisa Linke, Alexandro Rodríguez-Rojas, Jens Rolff and Seulkee Yang for advice and/or technical support. We thank Jens Rolff for feedback on an earlier draft of the manuscript.

## Data Accessibility

Data will be made publicly available upon acceptance of this article.

## Supporting Information

**S1 Fig. Illustration of the predictions for how clearance is expected to change with A. per parasite pathogenicity (PPP) and B. exploitation.**

**S2 Fig. Per parasite pathogenicity using different smoothing parameter values to estimate the maximum hazard.**

**S3 Fig. Effect of exploitation (bacterial load) on clearance for different values of the smoothing parameter.**

**S1 File. Statistical methods and results for the comparison between the proportions of live and dead uninfected flies.**

**S1 Table. Tukey multiple comparisons between bacterial species for differences in virulence.**

