## Supplementary Material for "Decomposing virulence to understand bacterial clearance in persistent infections"

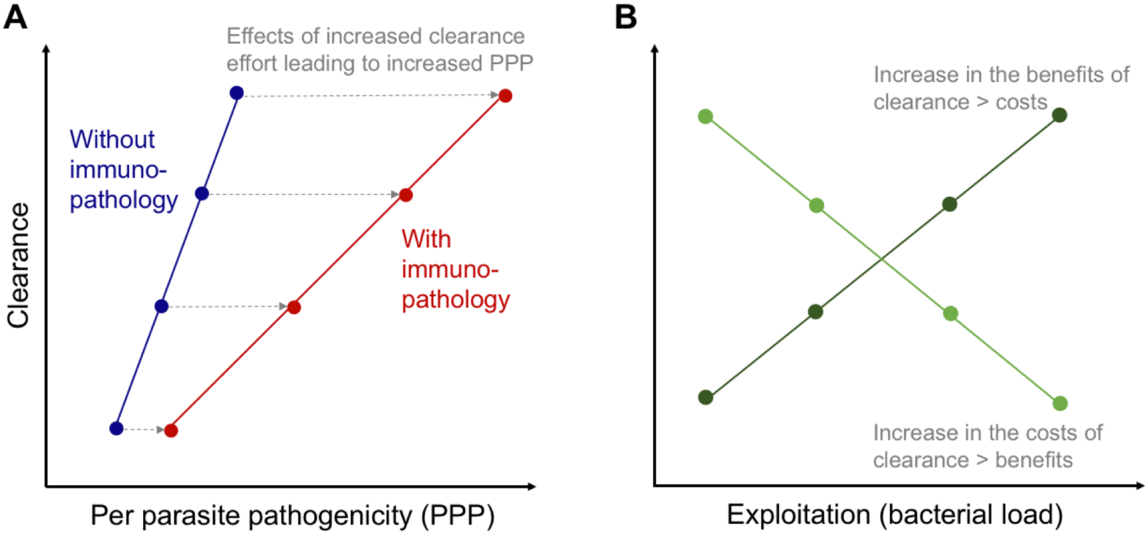


**S1 Fig. Illustration of the predictions for how clearance is expected to change with A. per parasite pathogenicity (PPP) and B. exploitation**. **A**. Increasing PPP should lead to increasing benefits of clearance, while having no or a negligible effect on the costs of clearance. Therefore, with increasing PPP, clearance should increase (dark blue line). Immunopathological effects can be expected to modify this pattern (*e.g.*, red line), but they should not reverse the relationship between PPP and clearance. In addition to the fitness loss caused directly by the pathogen, immunopathological effects will further decrease the fitness of infected individuals and thus should result in a higher PPP. In addition, we can expect that increased clearance efforts lead to an increase in immunopathology. Accordingly, with an increase in clearance effort, so the measured PPP also increases, which results in a flatter relationship between PPP and clearance (red line compared to blue line). **B**. Exploitation is expected to lead to an increase in the costs and the benefits of clearance. Depending on whether benefits or costs increase faster, clearance should either increase (dark green line) or decrease (pale green line) with increasing exploitation.

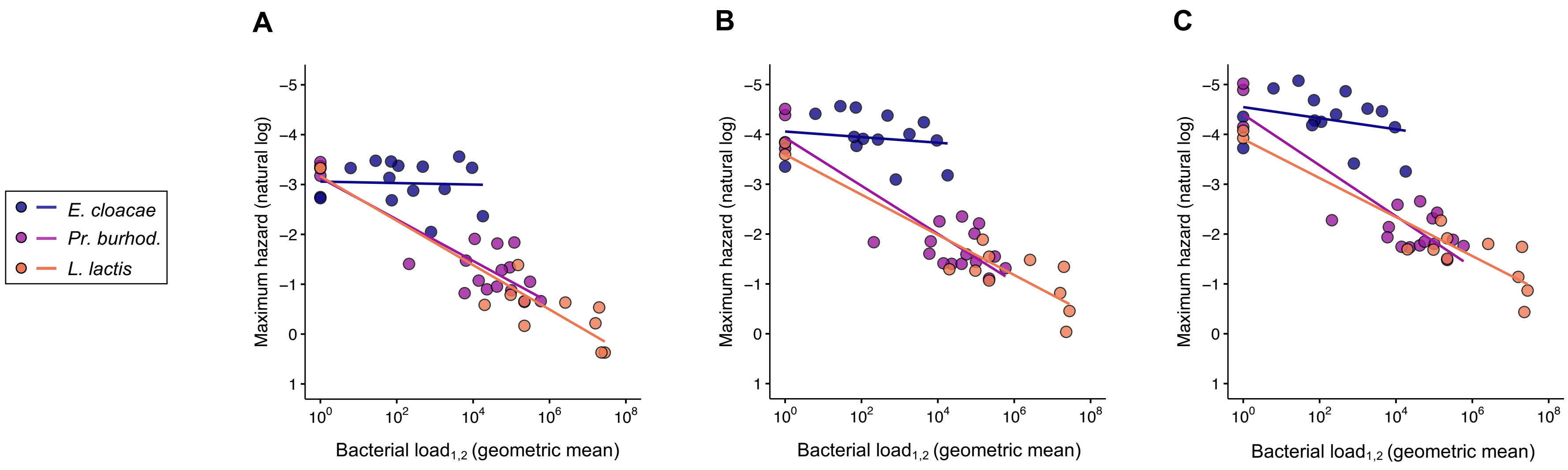


| | df | F | P | df | F | P | df | F | P |
|---|---|---|---|---|---|---|---|---|---|
| Bacterial species | 2,39 | 0.047 | 0.95 | 2,39 | 0.71 | 0.50 | 2,39 | 1.34 | 0.27 |
| Bacterial load | 1,39 | 0.027 | 0.87 | 1,39 | 0.34 | 0.56 | 1,39 | 1.30 | 0.26 |
| Bacterial species × bacterial load | 2,39 | 8.92 | **0.00065** | 2,39 | 7.29 | **0.0020** | 2,39 | 6.02 | **0.0053** |

**S2 Fig. Per parasite pathogenicity using different smoothing parameter values to estimate the maximum hazard.** Per parasite pathogenicity is given as the relationship between bacterial load and maximum hazard. The bacterial load data is the same as that given in Fig 3A, with the addition of the Ringer's treatment control. The maximum hazard data is estimated from survival data for the corresponding injection doses and experimental replicates. Maximum hazard is plotted as the inverse, such that the hazard (virulence) increases with proximity to the x-axis. The maximum hazard was estimated from time to death data using four different values (1, 2, 3 and 5) for the smoothing parameter, *b,* as specified using "bw.grid". Shown above are **A.** *b* = 1, **B.** *b* = 3, **C.** *b* = 5 *(b* = *2* is shown in Fig 3B). Coloured lines show the linear regressions. The corresponding statistical results are shown below each panel, where maximum hazard was the dependent variable. Statistically significant p-values are in bold.

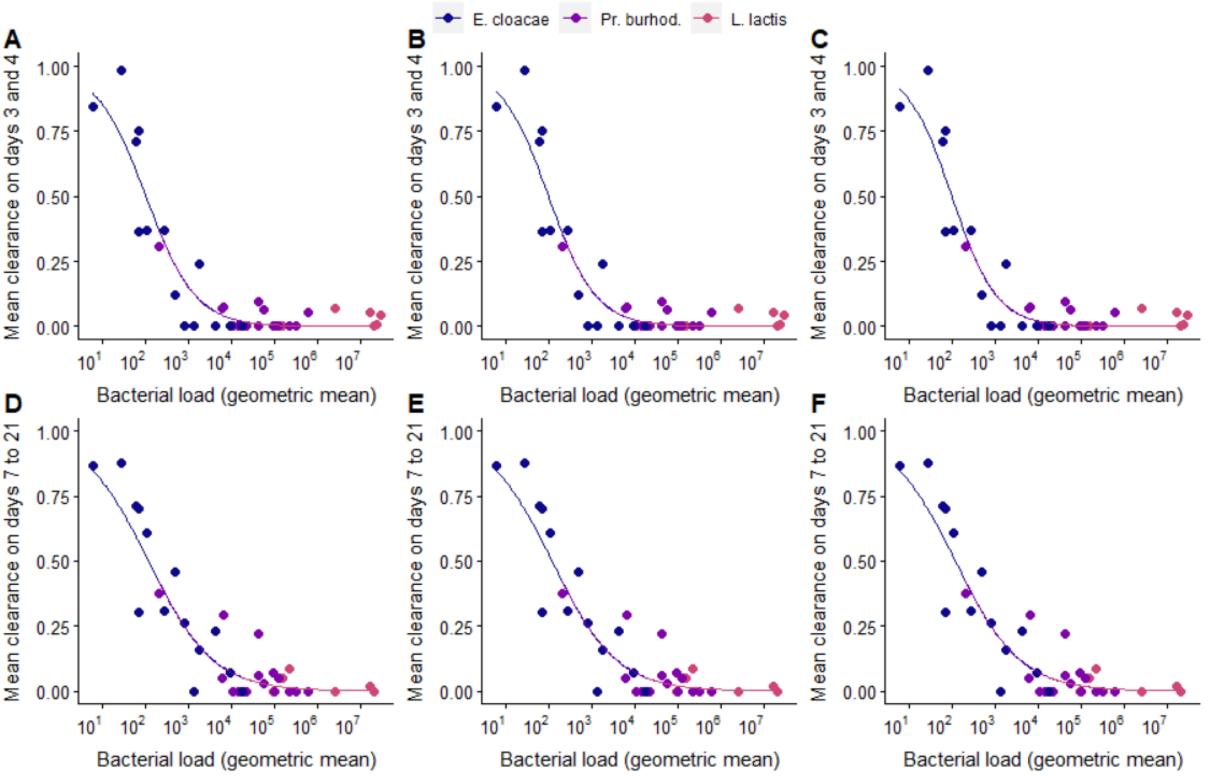


**S3 Fig.** Effect of exploitation (bacterial load) on clearance for different values of the smoothing parameter *b* in the estimation of maximum hazard that is used for the calculation of PPP: *b* = 1 (**A**, **D**), *b* = 3 (**B**, **E**), *b* = 5 (**C**, **F**). The geometric mean of bacterial load was calculated from days 1 and 2 post injection. Each data point is from one injection dose per bacteria, per experimental replicate, and gives the mean proportion of cleared infections on days three and four (**A**-**C**) and days 7 to 21 (**D**-**F**).

**S1 File. Statistical methods and results for the comparison between the proportions of live and dead uninfected flies.**

To test whether the proportion of live uninfected flies was a predictor of the proportion of dead uninfected flies, we separately summed up the numbers of uninfected and infected flies for each bacterial species and dose, giving us a total sample size of n = 20 (four species × five doses). For live and for dead homogenised flies we had a two-vector (proportion infected and proportion uninfected) response variable, which was bound into a single object using cbind. The predictor was live flies, and the response variable was dead flies, and it was analysed using a generalized linear model with family=quasibinomial.

Model 9: cbind(dead uninfected, dead infected) ~ cbind(live uninfected, live infected)

Despite variation in the time post infection at which live and dead flies were sampled, across bacterial species and doses and as expected, the proportion of living flies that cleared an infection was a predictor for the proportion of dead flies that cleared an infection (S4 Fig; LR = 7.11, df = 2,17, p = 0.0285). Most of the data points lie above, rather than on, the diagonal (S3 Fig) possibly because the dead flies were on average homogenised later on in the infection, giving time for more clearance to take place before being sampled.

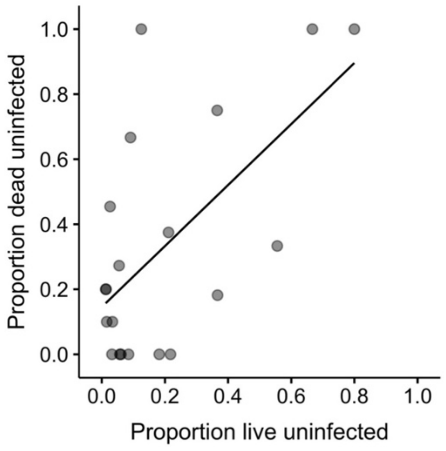


**S4 Fig. Proportion of live and dead flies that were uninfected across bacterial species and doses.** Each data point is the proportion for one bacterial species and dose. Darker circles are due to overlapping data points. The black line shows the linear regression.

**S1 Table. Tukey multiple comparisons between bacterial species for differences in virulence**. Virulence is measured as maximum hazard (model 1). Statistically significant comparisons are in bold.

| *Contrast* | *df* | *t* | *P* |
|---|---|---|---|
| *E. cloacae – L. lactis* | 55 | -17.23 | **< 0.0001** |
| *E. cloacae – Pr. burhodogranariea* | 55 | -13.42 | **< 0.0001** |
| *E. cloacae – Ps. entomophila* | 55 | -23.29 | **< 0.0001** |
| *L. lactis – Pr. burhodogranariea* | 55 | 3.88 | **0.0016** |
| *L. lactis – Ps. entomophila* | 55 | -6.17 | **< 0.0001** |
| *Pr. burhodogranariea – Ps. entomophila* | 55 | -10.04 | **< 0.0001** |
